# *Klebsiella huaxiensis* sp. nov., recovered from human urine

**DOI:** 10.1101/335075

**Authors:** Yiyi Hu, Li Wei, Yu Feng, Yi Xie, Zhiyong Zong

**Affiliations:** Center of Infectious Diseases, West China Hospital, Sichuan University, Chengdu, China; Division of Infectious Diseases, State Key Laboratory of Biotherapy, Chengdu, China; Center for Pathogen Research, West China Hospital, Sichuan University, Chengdu, China; Department of Infection Control, West China Hospital, Sichuan University, Chengdu, China; Laboratory of Clinical Microbiology, Department of Laboratory Medicine, West China Hospital, Sichuan University, Chengdu, China

**Keywords:** *Klebsiella huaxiensis*, *Klebsiella*, *Klebsiella oxytoca*, taxonomy, genome

## Abstract

A *Klebsiella* strain, WCHKl090001, was recovered from a human urine sample in China in 2017. Phylogenetic analysis based on *gyrA* and *rpoB* housekeeping genes revealed that the strain was distinct from any previously described species of the genus *Klebsiella* though it was clustered with the *Klebsiella oxytoca* phylogroup including *Klebsiella grimontii, Klebsiella michiganensis*, and *Klebsiella oxytoca*. The whole genome sequence of strain WCHKl090001 has an up to 87.18% average nucleotide identity with those of type strains of all known *Klebsiella* species. *In silico* DNA-DNA hybridization (isDDH) values between strain WCHKl090001 and type strains of all known *Klebsiella* species range from 22.3 to 35.2%. Strain WCHKl090001 could be distinguished from species of the *Klebsiella oxytoca* phylogroup by its negative Voges-Proskauer reaction. Genotypic and phenotypic characteristics from this study indicate that strain WCHKl090001 should be considered to represent a novel species of the genus *Klebsiella*, for which the name *Klebsiella huaxiensis* sp. nov. is proposed. The type strain is WCHKl090001^T^ (=GDMCC1.1379^T^ = CCTCC AB 2018106 ^T^).

---

*Klebsiella* is a genus of Gram-negative, non-spore-forming bacteria within the family *Enterobacteriaceae*. *Klebsiella* strains are widely distributed in nature and some *Klebsiella* species, in particular, *Klebsiella pneumoniae* is a common human and animal pathogen causing a variety of infections such as bacteremia, pneumonia, meningitis, urinary tract infection and intra-abdominal infection. In addition to the well-known pathogen *K. pneumoniae*, the genus of *Klebsiella* currently comprises *Klebsiella aerogenes* (also known as *Klebsiella mirabilis* and *Enterobacter aerogenes*) [1], *Klebsiella granulomatis* [2], *Klebsiella grimontii* [3], *Klebsiella michiganensis* [4], *Klebsiella oxytoca, Klebsiella quasipneumoniae* [5], *Klebsiella quasivariicola* [6] and *Klebsiella variicola* [7]. *Raoultella ornithinolytica, Raoultella planticola* and *Raoultella terrigena* were used to belong to the genus *Klebsiella* but have been transferred to the genus *Raoultella* [8]. During our clinical works, we found that a *Klebsiella* clinical strain, WCHKl090001, is distinct from all hitherto known species and therefore may represent a novel species of the genus *Klebsiella*.

Strain WCHKl090001 was recovered from the urine of a patient at West China Hospital of Sichuan University, Chengdu, China, in November 2017. The 16S rRNA gene sequence of strain WCHKl090001 was obtained by PCR using the universal primers 27F and 1492R [9] and Sanger sequencing. The nearly-complete 16S rRNA sequence of strain WCHKl090001 was closest (98.5% identity) to NBRC105695^T^, the type strain of *K. oxytoca*. However, the 16S rRNA sequence within *Klebsiella* is highly conserved and is unable to assign *Klebsiella* strains to the species level due to the limited phylogenetic resolution [3, 10, 11].

Strain WCHKl090001 was subjected to whole genome sequencing with 150 × coverage using the HiSeq X10 Sequencer (Illumina, San Diego, CA), which generated 1.49 Gb clean bases. Reads were trimmed using Trimmomatic [12] and were assembled to 149 contigs with a 53.3% GC content using SPAdes v3.11.1 [13]. Whole genome sequences are available for type strains of the genera *Klebsiella* and *Raoultella* except *K. granulomatis*, due to the fact that this species has not yet been cultured axenically [2]. The 383-bp internal sequence fragment of the *gyrA* gene (encoding DNA gyrase subunit A) and 501-bp of the *rpoB* gene (encoding RNA polymerase β-subunit) of type strains of the genera *Klebsiella* were retrieved from their whole genome sequences (Table 1). *R. ornithinolytica, R. planticola* and *R. terrigena* were also included in comparison as they were closely related to *Klebsiella* species. The concatenated sequences were aligned by MEGA 7.0 [14] to infer a maximum-likelihood tree. Strain WCHKl090001 is clustered with type strains of species of the *Klebsiella oxytoca* phylogroup including *K. grimontii, K. michiganensis* and *K. oxytoca* (Figure 1).

**Table 1.** ANI and isDDH values between strain WCHKl090001^T^ and the type strains of *Klebsiella* species.

| Species | Strain | Accession no. | ANI<br>(%) | DDH<br>(%) |
| --- | --- | --- | --- | --- |
| *K. aerogenes* | KCTC 2190<sup>T</sup> | CP002824 | 81.40 | 26.0 |
| *K. grimontii* | 06D021<sup>T</sup> | FZTC01000000 | 87.09 | 35.0 |
| *K. michiganensis* | H1g<sup>T</sup> | AYMI01000000 | 87.18 | 35.2 |
| *K. oxytoca* | NBRC105695<sup>T</sup> | BCZK01000000 | 86.74 | 33.8 |
| *K. pneumoniae* subsp. *ozaenae* | ATCC 11296<sup>T</sup> | CDJH01000000 | 81.41 | 25.6 |
| *K. pneumoniae* subsp. *pneumoniae* | ATCC 13883<sup>T</sup> | JOOW01000000 | 81.27 | 25.5 |
| *K. pneumoniae* subsp. *rhinoscleromatis* | ATCC 13884<sup>T</sup> | CDOT01000000 | 81.24 | 25.4 |
| *K. quasipneumoniae* subsp. *quasipneumoniae* | 01A030<sup>T</sup> | CCDF01000000 | 81.44 | 25.8 |
| *K. quasipneumoniae* subsp. *similipneumoniae* | 07A044<sup>T</sup> | CBZR01000000 | 81.48 | 25.6 |
| *K. quasivariicola* | KPN 1705<sup>T</sup> | CP022823 | 81.47 | 25.7 |
| *K. variicola* | DSM 15968<sup>T</sup> | CP010523 | 81.40 | 25.7 |
| *R. planticola* | B43<sup>T</sup> | BADH01000000 | 76.86 | 22.3 |
| *R. ornithinolytica* | ATCC 31898<sup>T</sup> | NC_021066 | 82.42 | 27.0 |
| *R. terrigena* | ATCC 33257<sup>T</sup> | LANE01000000 | 82.49 | 27.3 |

**Figure 1.**
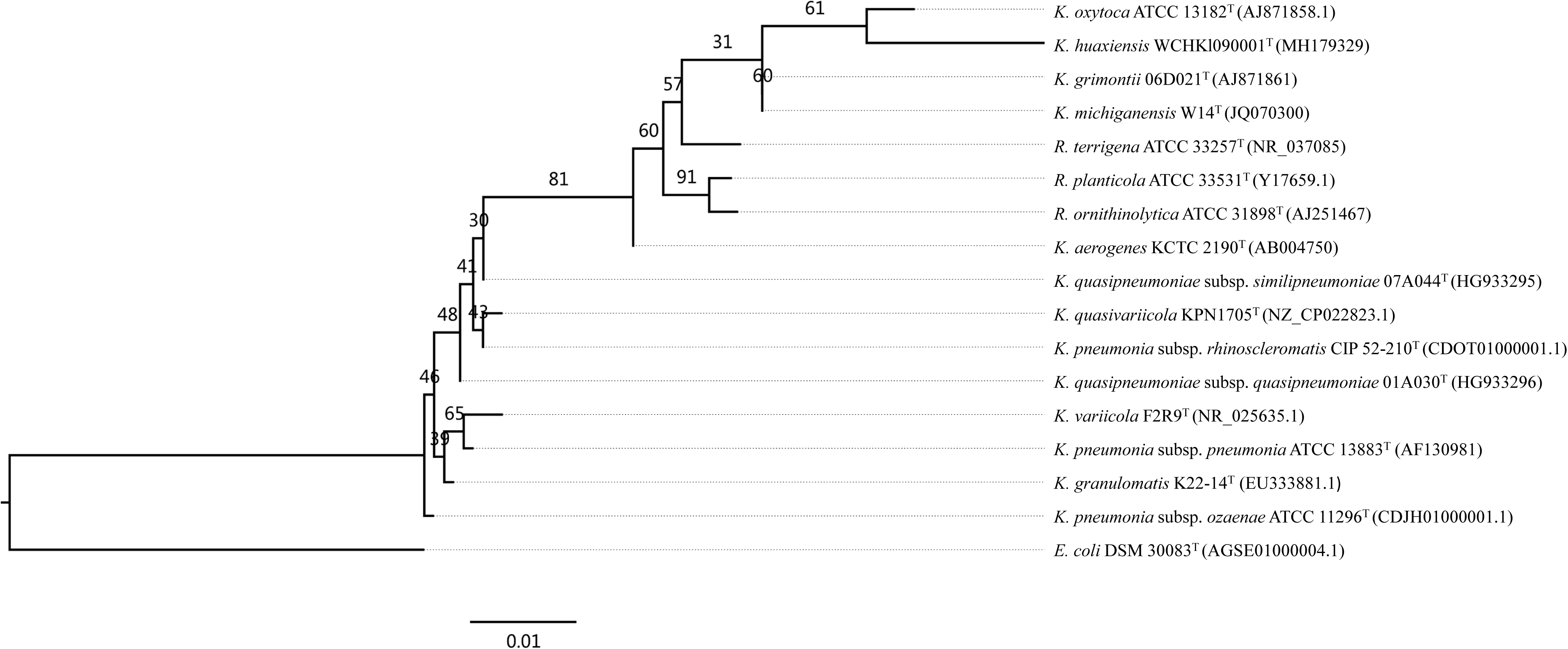
Neighbour-joining tree based on the concatenated sequences of the *gyrA* and *rpoB* genes of strain WCHKl090001^T^ and type strains of *Klebsiella* and *Raoultella* species. *E. coli* DSM30083^T^ (Accession no. AGSE01000004.1) was used as an outgroup. Bootstrap values > 50% (based on 1,000 resamplings) are shown. Bar, 0.005 substitutions per nucleotide position.

To further investigate the taxonomy of strain WCHKl090001, both the pair-wise average nucleotide identity (ANI) and *in silico* DNA-DNA hybridization (isDDH) between strain WCHKl090001 and type strains of the genera *Klebsiella* and *Raoultella* were determined. ANI was determined using the JSpecies web program based on BLAST [15]. Strain WCHKl090001 shared only 76.86 to 87.18% ANI with type strains of all known *Klebsiella* and *Raoultella* species (Table 1), which were well below the ≥95% ANI cutoff to define a bacterial species [16]. isDDH was performed using GGDC (formula 2) [17]. The isDDH relatedness between strain WCHKl090001 and any of the known *Klebsiella* and *Raoultella* species ranges from 22.3 to 35.2% (Table 1), much lower than the ≥70% cutoff to define a bacterial species. The ANI and isDDH analyses suggest that strain WCHKl090001 represents a new species of the genus *Klebsiella*.

Biochemical property of strain WCHKl090001 was determined using the API 20E kit and API 50CH kit according to the manufacturer’s instructions (bioMerieux, Lyon, France). All tests were carried out by incubating at 35 °C unless indicated otherwise. Biochemical characteristics of strain WCHKl090001 were compared with type strains of *Klebsiella* and *Raoultella* species. Strain WCHKl090001 was non-motile by microscopy. Strain WCHKl090001 was positive for indole, lysine decarboxylase, lactose, mannitol, and the ONPG test, reduced nitrate to nitrite but was negative for Voges-Proskauer test, malonate, urease and ornithine decarboxylase. The negative for Voges-Proskauer test could distinguish strain WCHKl090001 from other species of the *Klebsiella oxytoca* phylogroup.

Genotypic and phenotypic characteristics and the genome sequence of strain WCHKl090001 lend the support that the strain should be considered to represent a novel species of the genus *Klebsiella*, for which the name *Klebsiella huaxiensis* sp. nov. is proposed. The type strain is WCHKl090001^T^.

## Description of *Klebsiella huaxiensis* sp. nov

*Klebsiella huaxiensis* (hua.xi.en’sis. N.L. masc. adj. *huaxiensis* belonging to West China [Huaxi in Chinese] Hospital, Chengdu, Sichuan Province, China, where the type strain was recovered).

Cells are Gram-negative, non-motile, gas-producing, and capable of growing on media such as TSA (Oxoid, Hampshire, UK), LB agar, BHI agar and MH agar (all from Hopebio, Qingdao, China). Colonies on BHI agar after 24 h of incubation at 37 °C are light yellow, circular, smooth, convex, glistening, with entire margins.

*K. huaxiensis* belongs to the *Klebsiella oxytoca* phylogroup including *K. grimontii, K. michiganensis*, and *K. oxytoca*. The phenotypic characteristics of *K. huaxiensis* strain WCHKl090001^T^ are generally consistent with those for the *K. oxytoca* phylogroup. Positive for indole, lysine decarboxylase indole, lysine decarboxylase, lactose, mannitol, and the ONPG test, reduced nitrate to nitrite but negative for malonate, urease and ornithine decarboxylase. Negative reaction for Voges-Proskauer test could distinguish *K. huaxiensis* from other species of the *Klebsiella oxytoca* phylogroup. The G+C content is 53.3%.

The type strain is WCHKl090001^T^, recovered from a urine culture of a patient at West China Hospital of Sichuan University, Chengdu, China in November 2017. It is Voges-Proskauer test negative. The GenBank/EMBL/DDBJ accession numbers of the *gyrA, rpoB* and *rrs* (coding for 16S rRNA) genes are MH190069, MH190071 and MH179329, respectively. The genome sequence accession number is QAJT00000000.

Strain WCHKl090001 has been deposited into China Center for Type Culture Collection as CCTCC AB 2018106 and into Guangdong Microbiology Culture Center as GDMCC 1.1379.

